# Water Stress Induces Diel Patterns of Root Growth

**DOI:** 10.64898/2026.01.12.698942

**Authors:** Richard Nair, Martin Strube, Javier Pacheco-Labrador, Marion Schrumpf, Mirco Migliavacca

## Abstract

Growth and death of vegetation control both the amount of carbon held in living biomass and much of the flow between the atmosphere and soils, the largest terrestrial carbon reservoir. However, long-term field datasets on belowground growth dynamics are rare. Knowledge is lacking on how root growth patterns relate to the environment, fluctuating carbon and water availability, and internal plant physiological control, at both seasonal and diel scales. We imaged roots using automated minirhizotrons and interpreting root length density (RLD) automatically twice a day for nine months in a temperate grassland. From winter to autumn, we always observed some root growth in individual images, but with decreasing net RLD (indicative of net turnover) becoming predominant later in the season. Roots grew day-round except during a period of about one month in spring when root growth was only in daytime. This was a period without rain, during which soil moisture depleted rapidly, but not at its driest point. We discuss potential causes for both overall and short-term growth patterns, particularly the change in drivers of growth from internal storage to new photosynthates, or limits imposed by water availability - both soil moisture and rainfall - under shifting environmental conditions.

## INTRODUCTION

High-frequency measurement of ecosystem carbon (C) and water fluxes is now routine (Baldocchi, 2020), revealing patterns in land-atmosphere exchange, often due to combined variation in meteorology and partially coupled plant growth and turnover (e.g. Y. Liu & Wu, 2020; C. Wu et al., 2012). However, validation from whole-plant dynamics in-field is often opaque. While remote sensing provides above-ground information, at least 1/3 of plant biomass (Jackson *et al*., 1996; McCormack *et al*., 2015) and half of total productivity (Gherardi & Sala, 2020) is allocated belowground to ‘cryptic’ roots, mycorrhizal fungi, and exudates. When studied, root phenology is commonly asynchronous with leaves (Abramoff & Finzi, 2015), but measurements of roots at high frequency are also far less frequent. Overall, observational data on root growth dynamics on diel to seasonal scales is sparse compared to above-ground observations.

At seasonal scale, precipitation, temperature, and light control carbon and water availability, and soil nutrient turnover (e.g. Graham et al., 2003; Huete et al., 2006; Z. Wu et al., 2021), and ultimately plant phenology (e.g. Luo et al., 2020; Zelikova et al., 2015). Meteorological conditions and light availability also oscillate between day and night, influencing resource supply above-ground. Circadian rhythms help plants buffer these conditions, but activity still has diel patterns (Rozema *et al*., 1987; Walter *et al*., 2009; Poiré *et al*., 2010). Cell turgor limits initial leaf expansion (Pantin *et al*., 2011), often favouring nighttime (Rozema *et al*., 1987; Poiré *et al*., 2010), and similar patterns underly radial tree stem growth (Zweifel *et al*., 2021). Above-ground, turgor limits imposed by hydraulic conductance are a key growth driver.

Roots are located at opposite ends of the vascular system to leaves, so growth may relate differently to water and carbon (Ruts *et al*., 2012). They inhabit a dense, environmentally buffered, heterogeneous environment (Morris & Nair, 2025) and have different meristem mechanisms to shoots (Stahl & Simon, 2010). Daytime (Halter *et al*., 1996) and nighttime (Head, 1965) tree root growth has been reported in walk-in rhizotrons, but generally, roots tend to grow at night or around dawn (Ruts *et al*., 2012) and respond to environmental fluctuations (Bengough *et al*., 2011; Heinze *et al*., 2023; Ceolin *et al*., 2025). These data usually come from cuttings, isolated plants, short observations, or artificial conditions (Walter *et al*., 2002, 2009, (Yazdanbakhsh & Fisahn, 2009), whereas high frequency field measurements are limited (c.f. Belaud et al., 2024; Kuwabe & and Ohashi, 2023).

Here we used automated minirhizotron root cameras (Nair *et al*., 2022) to examine sub-daily fine root growth for 9 months in a temperate grassland. Over 250,000 images were analysed using a convolutional neural network (Smith *et al*., 2022) to track root length density (RLD) and rooting depth.

We aimed to answer the following questions:

Q1: **Under which environmental conditions do roots grow and die?** We predicted that we would observe the highest growth rates during warm periods when soil moisture was not limiting. We expected to see accelerating growth due to the cumulative effect of increased green biomass through spring, leading to increased C available to grow roots. We expected a seasonal cycle with net turnover later in the year.

Q2: **Is there a diel pattern of root growth, and if so, does this shift seasonally?** Following the evidence above, we expected night-time growth (if turgor limited) or no pattern (if it didn’t), without priors for seasonal changes beyond soil water availability.

## RESULTS

Net RLD growth was immediate when measurements began in February and continued until June, when it started to decline. Although the instruments matched each other in terms of gross patterns (Figure S3), the absolute values of RLD observed at each instrument microsite differed (Figure 1a). The mean rooting depth (MRD) deepened with a deepening soil moisture profile going into summer (Figure 1b, Figure 1d). The number of images with growing roots declined throughout the experiment (Figure S4). Minor recovery of MRD in September was driven by new roots in shallow soil layers (Figure S5).

**Figure 1.**
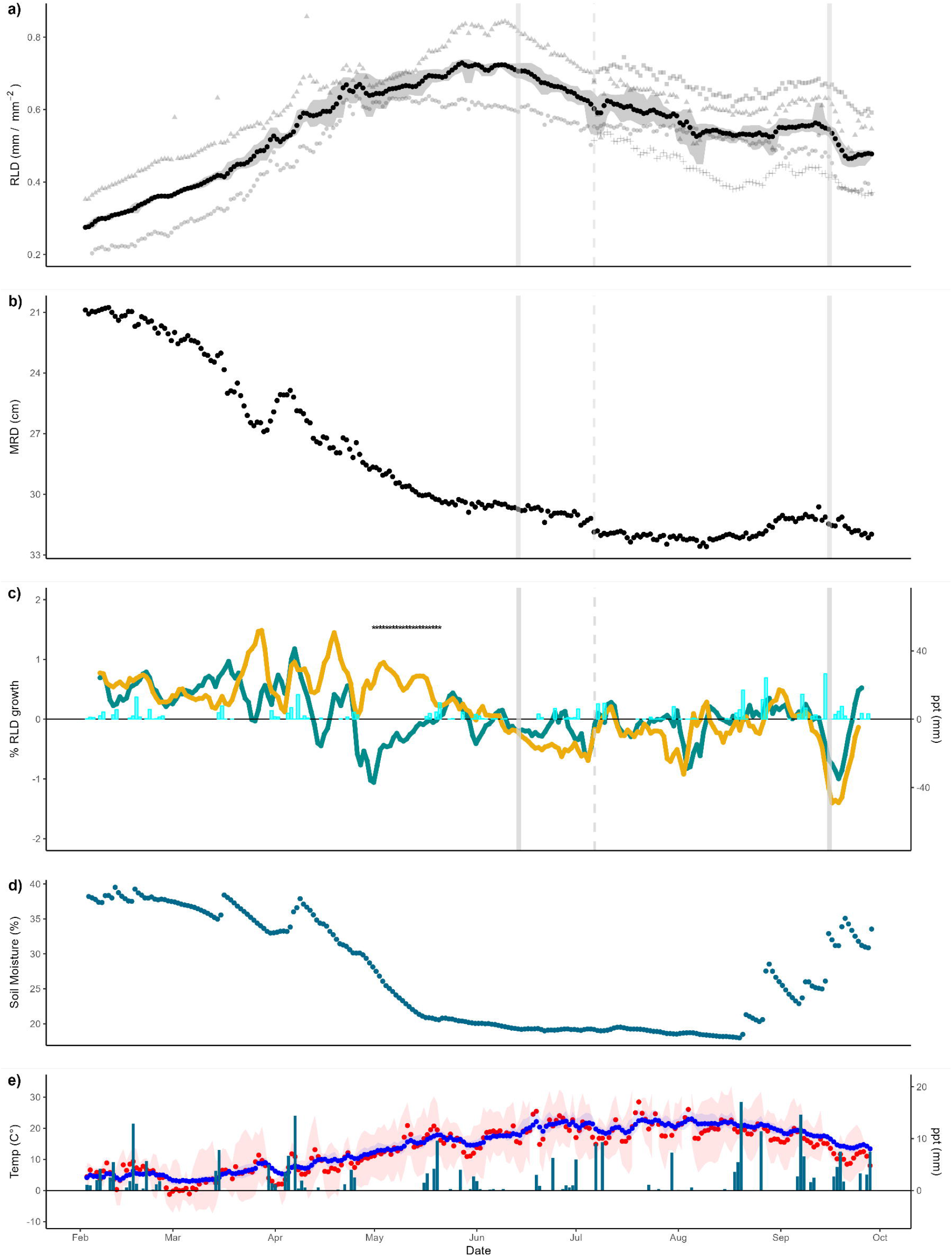
Root dynamics during the observation period. a) Mean root length density (RLD) dynamics between two instruments with two added in July (three day rolling mean, shapes indicate different instruments). Shaded area indicates the minimum and maximum observation within each five-day window. b) Mean rooting depth (MRD), the mean depth of roots observed in the minirhizotron. c) Sub-daily RLD growth. Throughout the observations, roots both grew (+ve growth) and turned over (-ve growth). Stars indicate periods with significantly different night-time growth than day-time growth (sign test). d) soil moisture as a % at measured 8 cm. e) rain, soil temperature (blue), and air temperature (red). The grey lines in the top three figures indicate (dashed) installation of new instruments, and (solid) first harvest and second mowing of above ground biomass removal in the management cycle.

We calculated daytime growth as the 3-day smoothed difference between the previous night and the current day, and vice-versa, for nighttime growth. Across the entire 40 cm depth, roots grew evenly between day and night until mid-April, when growth was predominantly during the day (Figure 1c). This returned to no difference by late May. Day-and-night growth was significantly different (sign test, P< 0.05) through most of May. No sustained or significant desynchrony was observed afterwards.

In April, there was a small but notable increase in rooting depth, which preceded decreasing soil moisture. Desynchrony between day and night occurred during this dry period, which was notable because was the longest precipitation gap in the entire timeseries (Figure 1c, Figure 1e).

## DISCUSSION

We investigated the timing of root growth in the field to identify 1) what environmental conditions roots grow and die, and 2) if there are diel patterns. We assumed that expansive growth is maximised at night/early morning, because during daytime, transpiration depletes water necessary for cell expansion (Walter *et al*., 2002). In lab-grown *Arabidopsis*, nighttime root elongation is under circadian control (Yazdanbakhsh *et al*., 2011) but imprints via hydrological conditioning (Caldeira *et al*., 2014). We found continuous root growth and only found diel desynchrony under late spring water deficits (Figure 1c). Here, more roots were produced during day than at night.

Growth could be limited by demand for resources, *source control* or other limits to use of these resources, *sink control* (Körner, 2015). Desynchrony could be due to 1) interruption or variation of C supply (source control) or 2) external temperature constraints, 3) a shift in internal partitioning or 4) interruption in water available for turgor. Explanations 2, 3, and 4 are sink controls. We discount shifts in endogeny because diurnal shifts only occurred at a single period, and our measurements were community scale. Overall, we interpret the interplay between carbon and water stress determining short-term desynchrony, as follows:

### Did carbon fixation (source control) control growth?

Net RLD increased steadily until May (Figure 1a), and some roots grew throughout the year even when net RLD declined (S3). Early in the growing season, C fixation was likely low, yet the trajectory of long-term RLD increase February-May was almost linear, suggesting no C limitation. Roots may respond slower to diurnal or weather-related shifts in C fixation than leaves due to C transport in the phloem, which operates via pressure-driven bulk flow and osmotic gradient regulation, (Münch, 1930; Mencuccini & Hölttä, 2010; Jensen *et al*., 2016). Transport time constraints are consistent with observed lags between photosynthesis and soil respiration in grassland ecosystems (Bahn *et al*., 2009). In such short-stature systems, leaf to root tip distance is short. Phloem sugar concentrations vary diurnally (Jensen *et al*., 2016), but with no consistent diurnal RLD change for most of our grassland timeseries, either pressure flow equilibrates supply and/or photosynthates were always sufficient. Either way, diurnal growth did not appear to be limited by C supply.

### Did temperature control growth?

Aboveground growth in temperate grasses becomes negligible below 5°C (Peacock, 1975; Wingler & Hennessy, 2016), but photosynthesis continues at lower temperatures (Körner, 2008) and leaf expansion is phytohormone regulated (Wingler & Hennessy, 2016). Soils are buffered from temperature and may be warmer than air; this was observed in February (Figure 1e), which may have allowed root growth despite stalled leaf production. RLD increased as soon as observations began (Figure 1a), initially mostly close to the surface (Figure 1b), followed by deepening roots as the season progressed. Summer mean daily air temperatures, rarely exceeding 25°C (optimal for temperate plants (Hatfield & Prueger, 2015)), were steadily increasing during the desynchrony period, suggesting temperature did not limit growth or drive the pattern.

### Did we miss carbon assignment to exudates or mycorrhizal fungi?

Diel asynchrony in root observations could potentially be due to variations in exudate or mycorrhizal components (i.e. total C remains the same, relative assignment different). Although automated minirhizotrons have observed mycorrhizal fungi (Allen & Kitajima, 2013; Defrenne *et al*., 2021), our systems were not designed for fungal hyphal resolution. Minirhizotron resolution usually trades off against field of view; high ‘zoom’ increases imaging time. Relevant fungal structures are further probably limited to rhizomorphs in ectomycorrhizal system (grasslands are dominated by arbuscular mycorrhizal fungi (Brundrett, 1991)) with less obvious effect on root phenotype. In any case, total C allocation to mycorrhizal fungi is around 1-13 % (Hawkins *et al*., 2023). In grasslands, most photo-assimilates also remain in plants in the short term (Ostle *et al*., 2007). From lab experiments, exudate patterns are also probably not diurnal (Badri *et al*., 2010), and we could not find diel patterns in mycorrhizal exchange. Complete belowground ^13^C traces through to efflux CO_2_ also do not reveal diurnal patterns (Liu *et al*., 2021). Thus, there are not likely alternate cryptic belowground sinks, which vary on diel or seasonal scales.

### Water and Turgor limited growth

Overall, seasonal patterns observed in this study matched root observations in similar ecosystems (Edwards *et al*., 2004; Steinaker & Wilson, 2008). Desynchrony (Figure 1c) occurred prior to a RLD decline over summer (Figure 1a). During this period, soil moisture (Figure 1e) and MRD (Figure 1b) decreased. Thus, the ecosystem was in transition from abundant to scarce soil water availability. Pasture grasses are susceptible to hydraulic failure and follow embolism avoidance strategies (Jacob *et al*., 2022). Leaf expansion is tightly coupled to soil water potential, but root growth is less so (Majerus, 1975), perhaps because of root proximity to the source of water . Because root extension may supply water, but leaves extends the transpiration stream, root growth may continue while leaf growth stalls.

Photosynthesis likely persisted under mild soil drought conditions, so there was fresh C available. Sentinel-2 Normalized Difference Vegetation Index (NDVI) remained stable during this desynchrony (Figure S6). Thus, daytime root growth could reflect diverted C from non-growing leaves, while nighttime was limited by water being used to refill depleted xylems. The plants may no longer have been able to reach turgor for root cell growth as the xylem refilled. Supporting this, MRD deepened before desynchrony, ahead of soil moisture decline, perhaps searching for additional water driven by leaf stress (Figure 1b, Figure 1d). The absence of rainfall during this period (Figure 1d) may have further shifted functional activity, with root building influenced by increasing leaf dryness and xylem vulnerability, even before soil moisture levels were fully depleted.

### Wider Implications

Growth timing matters for carbon budgets. In soil, respiration can be partitioned into root and heterotroph sources (Högberg *et al*., 2009; Taylor *et al*., 2015; Savage *et al*., 2018; Sun *et al*., 2019), or into growth, maintenance, and redox balancing components at plant level (De Vries *et al*., 1974; Johnson, 1990; Hirano *et al*., 2023). However, such partitioning requires assumptions, since data from undisturbed systems is scarce (Hirano *et al*., 2023). Respiration temperature sensitivity is usually modelled using Q_10_ enzyme kinetics, applied in ecosystem models (Meyer *et al*., 2018), partitioning of eddy covariance data (e.g. Reichstein *et al*., 2005), and interpreting processes such as acclimation and adaptation (King *et al*., 2006) plus other physiological insights. However, growth and maintenance have a different response to temperature, being driven by substrate sink/source limits and rates of reaction, respectively (De Vries *et al*., 1974). If their timing diverges, daily or long-term average Q_10_ may not reflect underlying processes.

Other evidence supports this view. Root temperature does not affect growth rates or respiration coefficients (Szaniawski & Kielkiewicz, 1982) but does affect total respiration via substrate supply (Atkin *et al*., 2000; Thurgood *et al*., 2014). Ecosystem respiration mostly comes from belowground (Raich & Schlesinger, 1992), with roots contributing 42 ± 18% (Jian et al., 2022). Root respiration has the highest Q_10_ of numerous factors contributing to soil CO_2_ fluxes (Boone *et al*., 1998), so our finding that root growth timing can switch between continuous and diel patterns under stress indicates such ecosystem generalizations may not hold consistently.

Similarly, models used to predict climate change are parameterised at half-hour scales for fluxes but fit to biomass over months to years. Fine roots are often simply prescribed via fixed ratios, thresholds, or optimality assumptions (e.g., Clark et al., 2011, Thum et al., 2019), without firm validation. Here we show diel desynchrony under environmental stress. Further work could link to above-ground dynamics and be used for model improvement, allowing teasing out source-sink controls and improving predictive capacity of coupled biogeochemical cycles. Understanding overall ecosystem activity may depend on what roots are doing and when they are doing it.

## MATERIALS AND METHODS

### Field Installation

The study was conducted at a grassland on the floodplain of the river Saale in Germany 50°55⍰N, 11°35⍰E. The site is a loamy Eutric fluvisol and hosts the Jena biodiversity experiment (Roscher *et al*., 2004; Weisser *et al*., 2017). We set up automated minirhizotron instruments in two ancillary plots without manipulation. The plots were unmanaged except for two cuts during the growing season when the above ground biomass was manually clipped above the instruments corresponding to mowing of the wider experiment.

We used an minirhizotron instrument design to capture images autonomously (Nair *et al*., 2022). In brief, this system took a series of images around a 1 m (length) x 10 cm (external diameter) observatory made of plexiglass. Roots grew around the observatory and were visible in the images. Image resolution was 25 μm per pixel, which covered the root diameters expected at the site. We installed observatories in September 2021 at 40 degrees to the horizontal, for a vertical span of ∼ 40 cm and a horizontal span of ∼ 70 cm, following best practice (Box *et al*., 1989; Johnson *et al*., 2001; Strand *et al*., 2018; Freschet *et al*., 2021) to ensure a tight fit between soil and tube. The length of ‘rest’ time before observations varies in the literature; here it was five months. Issues like soil-tube gaps and compaction may influence root behaviour (Bengough *et al*., 2006) and even follow hydrological changes in soil properties. As our rest period was over winter, it is likely that there was minimal root growth but time for soil expansion in the wetter part of the year.

We deployed two automatic minirhizotrons on 1 Feb 2022 and added a further two in Jul 2022. The instruments were in two clusters but because of the short depth of field of minirhizotron imagery (Johnson *et al*., 2001), we treated each minirhizotron as independent. The instruments ran on mains power and sampled at 01:00, 09:00, 13:00, and 21:00 UTC. At each sampling interval, the instrument imaged the entire observatory surface in 112 images over approximately 35 minutes. Lights, provided by a ring of LEDs angled away from the field of view, were switched on at image capture and off immediately afterwards. Images were saved in a .jpeg format and retrieved via exchange of an SD card on site. The instruments ran on mains power except on a few occasions where they failed to start; these missing data were gap filled as detailed later. Here we show data until 29 September 2022. This period covered the main growing season, with a gradual progression from winter conditions (air temperature min -7°C) through summer (air max. of 39 °C) and early autumn. Soil temperature was more buffered and at 8 cm depth reached minima of 1.6 °C. Even at the beginning of this dataset (the first 6 weeks are covered coarsely in Nair *et al*., 2022), root growth could be observed in sequential images. We could observe the disappearance of roots from images, indicating turnover or herbivory. Over the period of data collection there were periods with clear artefacts in the images, such as condensation and soil fauna, which we were mostly successfully able to train the Convolutional Neural Network (CNN) ignore in segmentations.

### Image Processing

Field minirhizotron imagery is still difficult to process automatically and while much progress has been made in using neural networks for root images in general this is largely in simple agricultural or mesocosm settings and designed on sparse timepoints (e.g. Gillert *et al*., 2021; Smith *et al*., 2022)) which are effective for sparse timepoints but had potentially high noise between sequential images. We smoothed the data as described later to offset this. Further development of sequential models could improve the interpretation of dense timeseries.

We processed every image collected using the same workflow (Figure 2), processing 268972 images over 2480 individual sampling cycles over four instruments). The size of this dataset meant we had to limit training data to a small fraction of the total image set, despite using a corrective annotation method, *rootpainter* (Smith *et al*., 2022) that allows a large throughput in training. Workflows based on *rootpainter* have been successfully used to interpret root images in both mesocosm (Alonso-Crespo *et al*., 2021) and field sites (Bauer *et al*., 2021; Nair *et al*., 2022). At this site, a cross-validated R^2^ of 88 % against was achieved image-level manual markup in the first six weeks of the experiment (Nair *et al*., 2022). Our philosophy for the interpretation of automatic instruments is to maximise *overall* consistency through time and at the scale of whole images than strict pixel-correct segmentation. Hence a declining value on our graphs is not disappearance of individual roots, but declining segmented area across all images.

**Figure 2.**
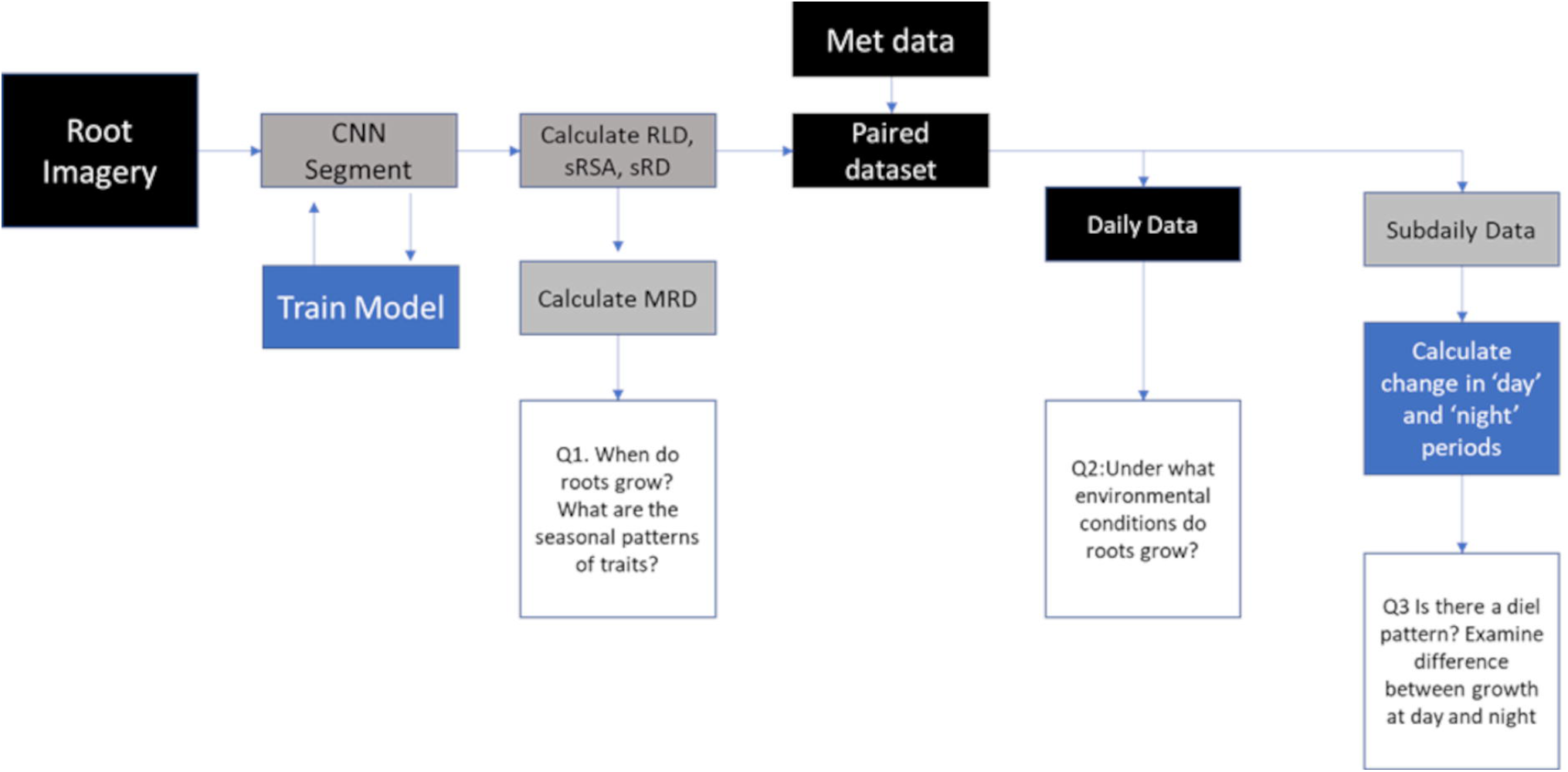
Summary of our workflow. Black boxes indicate data, grey boxes automated processing, and blue boxes automated processing where parameter choice was meaningful.

We excluded images on the top or bottom plane of the minirhizotron and those immediately at the soil surface, leaving 76 images per instrument per observation. We trained the model specifically on this site using a dataset consisting of 25 randomly selected images per instrument-week of the experiment, which were used to train in a random order. This dataset was fully generated after data collection was complete (i.e., not the same model as we used in Nair *et al*., (2022)) so there was not a bias in training the model from images in the sequence they were collected nor bias towards earlier images in the training. We trained for 350 random images at which point the f1 score indicated that the model was not improving. We repeated the complete training on an independent random set of 350 images and compared segmentations on 500 unseen images to validate that our training was reproducible (Figure S1). Overall, the two segmentations had a correlation coefficient r = 0.96.

Following segmentation, we processed binary pixel maps of the images to extract basic trait data, mostly in Ppython 3 using the skikit-image package. We followed the same protocol as (Nair *et al*., 2022) with the addition of an additional property, mean rooting depth (MRD). This was done as follows: first, we removed non-contiguous segmented areas of less than 0.5 mm^2^ to limit the influence of time-variable segmentation artefacts. We skeletonized the remaining segmented area to a single pixel-wide line, preserving original connectivity using medial axis segmentation and pruned all skeleton segments with a length of less than 10 pixels. Short segments were predominantly artefacts of the segmentation procedure and contributed little individually to the total root length. We counted all pixels in the remaining skeleton; we consider this root length. It contains a small artefact due to not adjusting for diagonal connections, but because roots are mostly linear in the images - unlike scanned root samples from soil core extractions - we expected this effect to be minimal. RLD (mm^2^ cm^-2^) is root length / image area. We calculated the *mean rooting* depth per instrument:timestep in R. As the minirhizotron observations were not evenly distributed in the vertical profile we fit a loess regression to generate the best fit smooth curve over the depth profile and predicted a RLD value at even 0.5 mm steps between 0 and 50 cm depth. We then proceeded through the predicted root length to find the depth at which 50% of the total root length form the model had been reached, which we treated as MRD (cm).

### Gap Filling

The image archive had a few gaps as on rare occasions the instruments did not start when scheduled. There were also instances when image artefacts occurred in the colour balance or brightness of the images due to data transfer from the camera, or power issues with the Light Emitted Diode (LED) lighting. We filtered out these occurrences automatically based on image colour statistics as these issues could affect the segmentation result. We also gap filled data points which were more than 25 % larger or smaller than the mean of the two observations before and after it; this accounted for short term anomalies such as soil animals which may obscure roots or be segmented as roots in a single image in a time series. Overall these data gaps affected 4% of images across the whole dataset. Having removed these data, we then gap filled the data for each trait via linear interpolation for a maximum gap size of 10 sequential observations. In practice, this meant an observation (real or gap filled) was available for every potential data point. Finally, in creating the RLD and MRD curves shown, we smoothed the data with a 3-day rolling average.

#### Meteorological and Remote Sensing Data

We used data from the MPI BGC Jena ‘Saaleaue’ meteorological station, located approximately 150 m from the root observations. The station provided us with air temperature, soil moisture (SM) content, and precipitation at the site level not paired to any instrument. For air temperature and moisture, we took the average of all sensors between 5 and 40 cm (the maximum depth of the minirhizotron). For precipitation, we averaged three rain gauges. The weather station data was available on 30 minute resolution, and further averaged to a daily averages to pair with daily averages growth rates from the minirhizotrons and calculated a further lagged 5 day (air temperature) and 14 day average (all meteorological variables except Rain and SM, which we left out because of their tendency to have strong effects on short timescales).

We lacked a dedicated paired above-ground canopy dynamics assessment paired to the minirhizotrons instruments. To assess whether the canopy was growing, we used Sentinel 2 imagery of the Jena Experiment site at 10 m spatial resolution, and computed Normalised Difference vegetation Index (NDVI) product with a cut-out of the Jena Experiment site for gross patterns of biomass above ground in our herbaceous system. There were 49 overpasses in the period of the experiment, of which 30 were obscured by clouds. With these removed from the dataset, we fit a LOESS regression across the remaining data for a smooth to produce a smooth curve that was used to interpret changes in the more frequent belowground data (Figure S6).

### Sub-daily Patterns

Minirhizotron data may be subject to variation on sub-daily timescales which are not growth patterns but real sub-daily patterns of cell turgor (Huck *et al*., 1970) or artefacts in neural network segmentation due to root or soil appearance (Nair *et al*., 2022). We wanted to discount these as much as possible to examine actual sub-daily growth. To do this, we started on the raw data (without the smoothing previously mentioned), observing that the biggest differences were between observations at midday and midnight – midday tended to have slightly higher RLD than midnight (Figure S2). This cycle was almost at inflection at 7 and 19 UTC without a significant difference in observed mean RLD between instruments when grouped by calendar week (Supplementary Table S2.1)). After we found the desynchronous behaviour, we re-ran this analysis, splitting the dataset into three groups: between the periods before, during, and after the desynchrony, and found significant effects of hour only in this period. Details in this post-hoc analysis are in Supplementary Table S2.2.

We hence treated the period between 7 and 19 UTC as ‘day’ and from 19 and to 7 as ‘night’. We smoothed the date across the time axis by applying a 3-days rolling average between day and night periods and calculated daytime growth as the difference between the previous NIGHT and the current DAY, and vice versa for nighttime growth.

To examine the trends in timing of growth over time we aggregated the DAY/NIGHT data arbitrarily into calendar week bins, each containing 7 observations of DAY and 7 of night from each instrument. We treated these as independent. We tested if day and night were significantly different for every sequential day-night and night-day pair of image sets within each calendar week using sign tests, a non-parametric method to test the median of two distributions.

## ACKNOWLEDGEMENTS

The authors are grateful for the Jena Experiment for hosting this study.

We also acknowledge technical help in minirhizotron development and maintenance from Agnes Fastnacht, Kerstin Hippler, Nadine Hempel and Tarek El-Madany.

## AUTHOR CONTRIBUTION

RN: Richard Nair, MS1: Martin Strube, JPL: Javier Pacheco-Labrador, MS2: Marion Schrumpf, MM: Mirco Migliavacca

Conceptualization: RN, MM. Methodology: RN, MS1, MS2, JPL. Investigation: RN. Data Curation, RN, Formal Analysis, RN, Software: RN. Writing – original draft: RN, Writing – review & editing: MS2, MM, JPL

## DATA AVAILABILITY

## Notes

### Competing Interest Statement

The authors have declared no competing interest.

### Summary of Updates

corrected some author affiliations which were incorrect

### Funding

- Research Ireland: URF\R1\231376

