## Supplementary Material for "Water Stress Induces Diel Patterns of Root Growth" (from another version of this preprint: version 1, 2026-01-12)

### Model Training

We could only train on a tiny fraction of the dataset, so we validated the training of our RootPainter model by training it twice on two randomly sampled training datasets. Overall the two models did not completely agree, but had a correlation coefficient of 0.96. Variation in the model is likely due to both images used to train and ‘noise’ in the input training data produced by the human annotator.

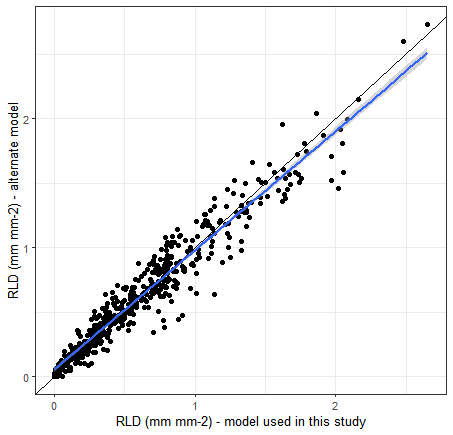

**Figure S1**. Comparison of two independently trained models on 500 random images. The segmentations had a correlation coefficient of 0.96.

### Identification of day and night points

We identified the 7 and 19 UTC periods through a visual inspection of the dataset (Figure S1.1). We smoothed the data on a 3 day window. This limited the effect of extremely short term errors due to automated segmentation (e.g. condensation, soil animals) while preserving general short term trends sustained over multiple days.

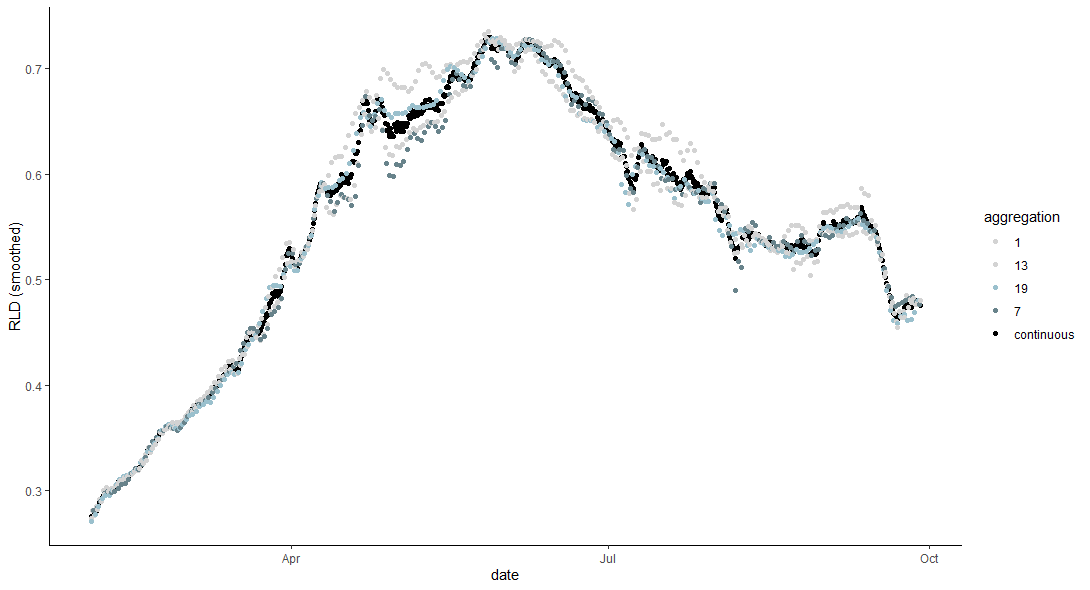

**Figure S2.** Comparison of RLD (3 day smooth) using the continuous, four times daily dataset, and when grouped into individual hour of observation. The RLD was consistently higher at the 13 UTC observation than the 1 UTC observation, especially in the latter part of the data so we discarded these observations and focused on the observations close to dawn and dusk.

On these 7 and 19 UTC observations, we performed several linear models to assess if there was a systematic bias on daily timing across the dataset (e.g. was RLD different at the end of the day, or at the end of night). In each case the data were transformed to be as normal as possible using Tukey’s ladder of powers. We conducted one analysis on the entire dataset, from February to September, and one in three parts – before the observations of desynchrony found in the paper (February to April), during the main period indicated by the sign tests (May) and after this (June to September).

**TableS2.1**. Summary of linear model on the whole dataset. Only week had a significant effect on the mean growth rate since the last observation.

|  | df | F value | P value |
| --- | --- | --- | --- |
| week | 1 | 57.2923187 | *** |
| hour | 1 | 0.001555597 | - |
| week:hour | 1 | 0.1185836 | - |
| Residuals | 472 | NA | NA |

**Table S2.2**: Summary of significance levels (* = P < 0.05, ** = P < 0.01, *** = P < 0.001) of linear models, when split between a) the period prior to, b) the period containing apparent desynchrony (May), and c) the period after desyncrhony. Hour was only significant in model b)

|  | P (February-April) | P (May) | P (June-September) |
| --- | --- | --- | --- |
| week | *** | *** | *** |
| hour |  | *** |  |
| week:hour |  | * |  |

### Constant Root Growth

At all times in our observations, some roots were growing, even during the periods of net decline in RLD (Figure S3).

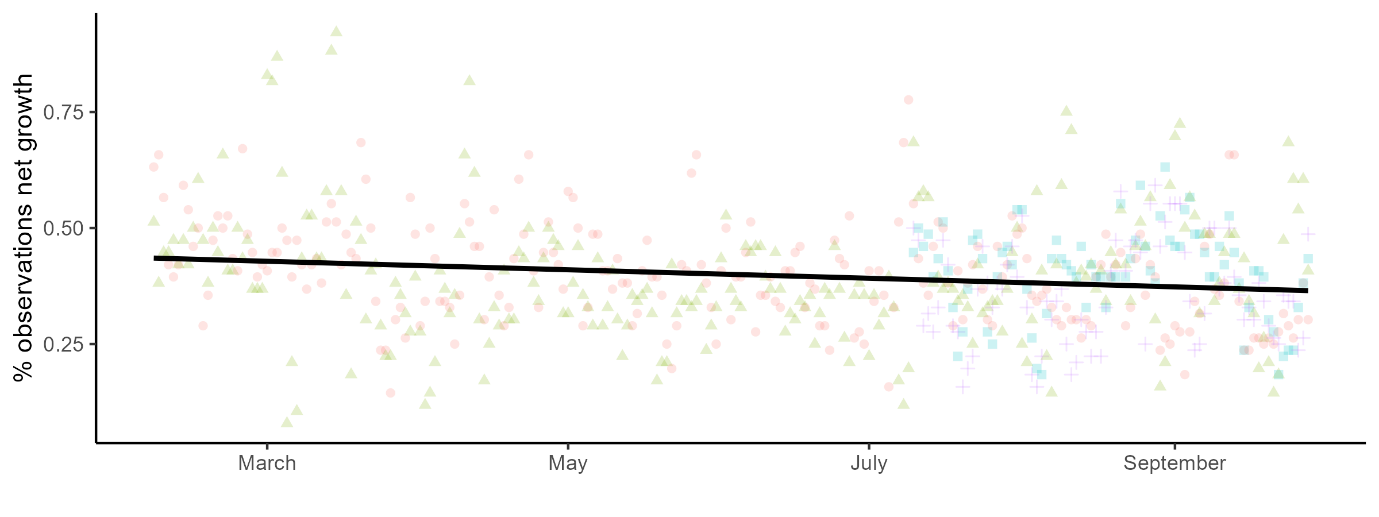

**Figure S3.** All timepoints of observation against the percentage of valid images with net growth (increase from previous timestep). At all times, there were some images with net growth observed.

### Depth Profile Distribution

Each minirhizotorns made observations across the root profile from the soil surface to 45 cm deep. These shifted during the seasons as evidenced by the MRD in (Figure 1). From the full profiles, we can see that rooting depth exceeded the minirhizotrons in summer (Figure S4). Our MRD estimates are thus likely an underestimate.

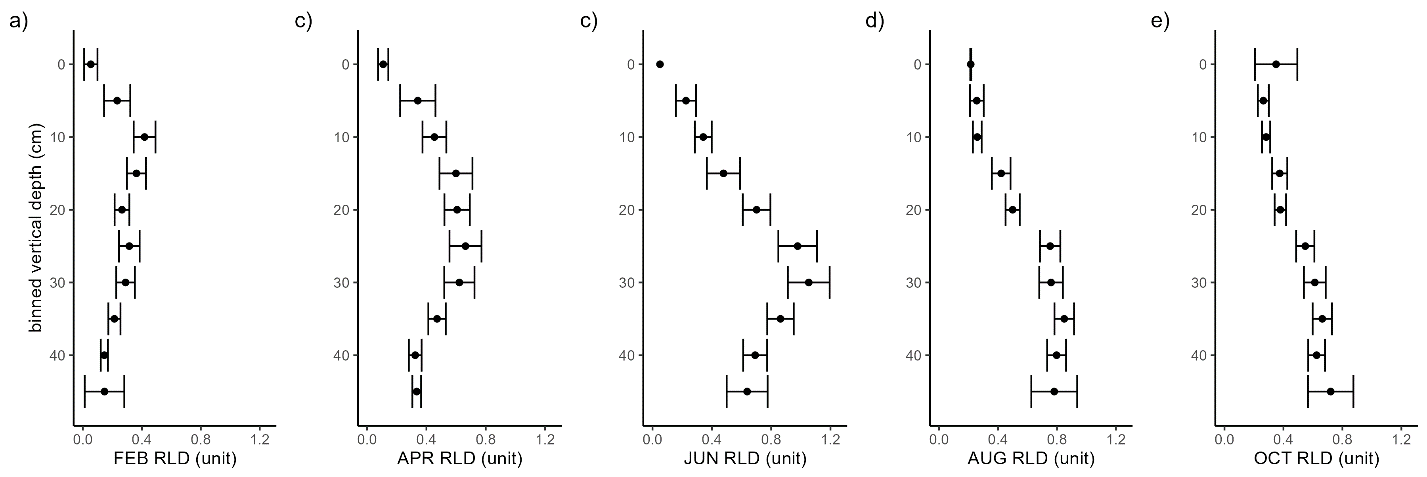
 **Figure S4.** Selected profiles of RLD in five months of 2023. The rooting profile shifted to deeper soil layers as the year went on but showed signs of recovery in shallow soil by October (Fig 2e) . We did not capture the entire depth, evidenced by the non-zero RLD in panels b,c,d,e.

### Comparison of Above- and Below-ground Timeseries

In this study, made during ad-hoc during the COVID lockdowns, we did not plan for the observations described in this paper. Therefore we do not have a proximal above-ground dataset available at similar time resolution. We used Sentinel2 NDVI as a rough proxy for vegetation change (Figure S5). Many of the overpasses were obscured by cloud but from this index it appeared that the vegetation change aboveground was stationary during our period of desynchrony. Due to the dry, precipitation-free conditions this was also the period where we had the most cloud-free overpasses and thus the best data quality in the timeseries.

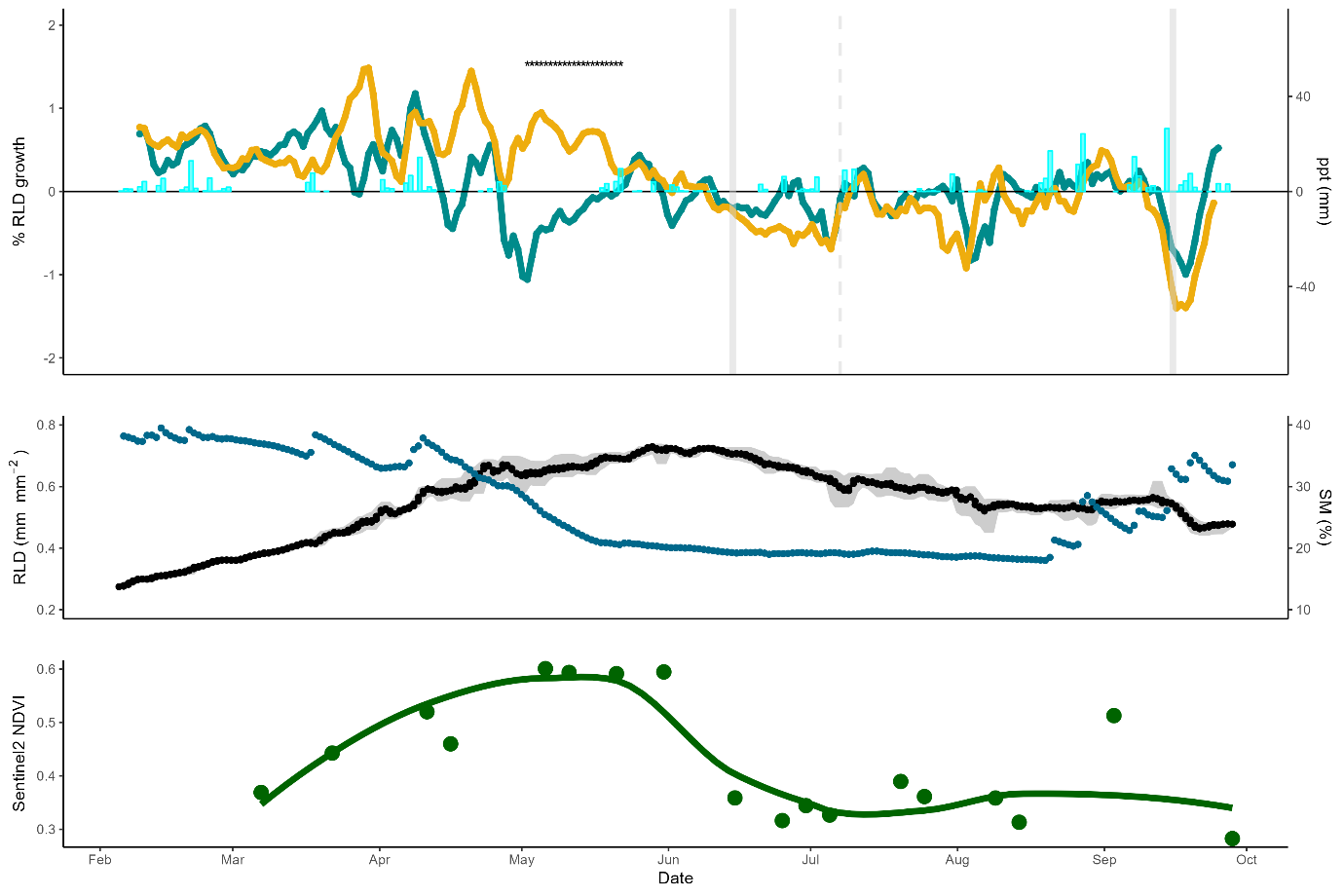

**Figure S4.** Comparison of above- and below-ground data. a) Growth desynchrony in day and night (as Figure 1, yellow and teal line) and precipitation (light blue bars) b) RLD (black) and soil moisture (blue), c) Sentinel-2 NDVI. Individual points are single cut-outs from a cloud-free overpass. The period of desynchrony indicated via sign tests in a) corresponds to the stationary period in panel c).
